# Harnessing the flexibility of neural networks to predict dynamic theoretical parameters underlying human choice behavior

**DOI:** 10.1101/2023.04.21.537666

**Authors:** Yoav Ger, Eliya Nachmani, Lior Wolf, Nitzan Shahar

## Abstract

Reinforcement learning (RL) models are used extensively to study human behavior. These rely on normative models of behavior and stress interpretability over predictive capabilities. More recently, neural network models have emerged as a descriptive modeling paradigm that is capable of high predictive power yet with limited interpretability. Here, we seek to augment the expressiveness of theoretical RL models with the high flexibility and predictive power of neural networks. We introduce a novel framework, which we term theoretical-RNN (t-RNN), whereby a recurrent neural network is trained to predict trial-by-trial behavior and to infer theoretical RL parameters using artificial data of RL agents performing a two-armed bandit task. In three studies, we then examined the use of our approach to dynamically predict unseen behavior along with time-varying theoretical RL parameters. We first validate our approach using synthetic data with known RL parameters. Next, as a proof-of-concept, we applied our framework to two independent datasets of humans performing the same task. In the first dataset, we describe differences in theoretical RL parameters dynamic among clinical psychiatric vs. healthy controls. In the second dataset, we show that the exploration strategies of humans varied dynamically in response to task phase and difficulty. For all analyses, we found better performance in the prediction of actions for t-RNN compared to the stationary maximum-likelihood RL method. We discuss the use of neural networks to facilitate the estimation of latent RL parameters underlying choice behavior.

**Author summary:** Currently, neural network models fitted directly to behavioral human data are thought to dramatically outperform theoretical computational models in terms of predictive accuracy. However, these networks do not provide a clear theoretical interpretation of the mechanisms underlying the observed behavior. Generating plausible theoretical explanations for observed human data is a major goal in computational neuroscience. Here, we provide a proof-of-concept for a novel method where a recurrent neural network (RNN) is trained on artificial data generated from a known theoretical model to predict both trial-by-trial actions and theoretical parameters. We then freeze the RNN weights and use it to predict both actions and theoretical parameters of empirical data. We first validate our approach using synthetic data where the theoretical parameters are known. We then show, using two empirical datasets, that our approach allows dynamic estimation of latent parameters while providing better action predictions compared to theoretical models fitted with a maximum-likelihood approach. This proof-of-concept suggests that neural networks can be trained to predict meaningful time-varying theoretical parameters.

## Introduction

A fundamental goal in neuroscience is to describe the underlying computations that animals and human agents might deploy when deliberating between actions. In pursuit of this goal, researchers have used reinforcement learning (RL) models which seek to describe behavior through a concise set of interpretable parameters that are believed to correspond to specific underlying processes (i.e., learning rate, exploration rate, etc.) [1–3]. While these models have generated important findings [4–6], they usually fit behavioral data poorly since they typically adopt a normative approach, focusing on how behavior should be rather than learning a generative model from the data itself [7, 8]. A more recent descriptive approach, in which neural networks are fitted directly to behavioral data, has been shown to exceed RL models in action prediction [9]. Neural networks are extremely flexible and can be used across a wide range of data variants and experimental environments [10, 11]. However, despite neural networks’ high predictive performance, they do not provide a clear theoretical interpretation, which is a major goal of brain and behavior research [12, 13]. Alternatively, neural networks could be employed to enhance and refine the parameter estimation for existing theoretical models, where the interpretation of parameters is more readily understandable.

Reinforcement learning models have proven effective in understanding and predicting decision-making behavior in both animals and humans [4, 14–16]. This method involves breaking down behavior into a parameterized computational model that can be fitted to individual choices [1, 2]. The individuals’ parameters are then viewed as low-dimensional representations of behavioral data, which provides insight into the latent cognitive configuration that might have generated the observed data. However, fitting RL models to behavioral observations can be challenging as it involves prolonged data collection in order to fit complicated and non-convex functions using iterative maximum-likelihood approximations [17]. To manage this complexity, constraints such as assuming stationary behavior [1] (i.e., fixed parameters) or rationality [8] (i.e., optimality) are imposed, eventually leading to lower predictive accuracy.

Recently, an alternative approach to studying decision-making has gained much attention, involving fitting neural network models directly to behavioral data [9, 18–20]. Unlike theoretical RL models that rely on assumptions about behavior, neural networks require minimal assumptions and can learn complex features from the data without human intervention. This data-driven approach has yielded high predictive accuracy on various behavioral tasks [9, 18]. Although some researchers have used neural network modeling beyond prediction to enhance our theoretical understanding of behavior [19, 21], the challenge of how to utilize the flexibility of neural networks to better interpret behavior remains open.

Here, we present a novel framework we term theoretical-RNN (henceforth, t-RNN), which aims to combine the strengths of RL models with the flexible and predictive power of neural networks. This is done by first simulating many artificial RL agents performing a two-armed bandit task, and then training an RNN using the trial-by-trial artificial data to predict both the agents’ theoretical RL parameters and future actions from observed action-reward history (see Fig. 1C). Moreover, the t-RNN recurrent structure allows it to estimate time-varying RL parameters, enabling it to track changes in behavior over time (see Fig. 1E).

**Fig 1.**
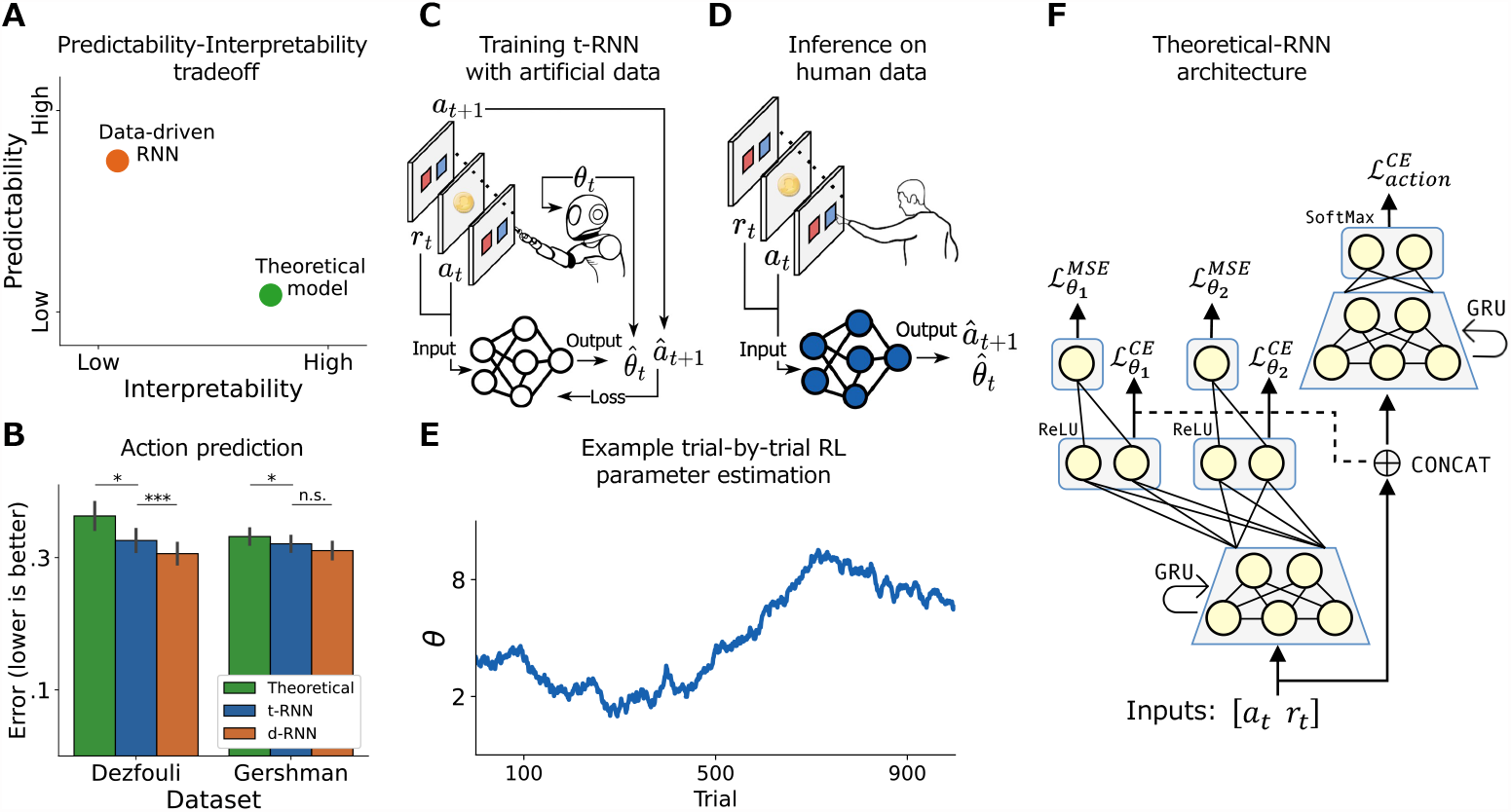
Summary schematic for theoretical-RNN. **(A)** Illustration of the predictability-interpretability trade-off plane. Theoretical models (green) are thought to be highly interpretable (x-axis), yet their predictive capability (y-axis) is often limited compared to data-driven RNN (d-RNN; orange). We suggest a new way to train a network (t-RNN) that allows dynamic estimation of theoretical RL parameters. **(B)** Empirical results for the analysis of two publicly available datasets where human individuals performed a two-armed bandit task [9, 22]. We present a comparison of action prediction made by d-RNN (orange), t-RNN (blue), and stationary theoretical modeling (green). Overall, we found that t-RNN was able to produce better predictions compared to stationary theoretical modeling fitted using a maximum-likelihood approach (y-axis illustrates error measured with binary cross-entropy; lower is better; black lines indicate s.e.m). **(C)** Illustration of t-RNN training procedure. We first simulated many artificial RL agents performing a two-armed bandit task with known RL parameters. We then familiarized t-RNN with the mapping between the action and reward (*a*_*t*_, *r*_*t*_) in trial *t*, to the action of the next trial *a*_*t*+1_ and the theoretical RL parameters in the current trial *θ*_*t*_. **(D)** Estimating humans RL parameters. Post-training, we froze the network weights and predicted the future action 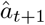, as well as estimated the dynamical RL parameters 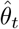, of humans performing a similar two-armed bandit task. **(E)** Illustration of t-RNN’s ability to estimate time-varying RL parameters, enabling it to track changes in behavior over time. **(F)** t-RNN architecture consists of several fully connected (FC) layers and gated recurrent units (GRU), such that the action prediction and RL parameters estimation are carried out with two different heads. Significant effect at * *p <* .05, *** *p <* .001; n.s., a nonsignificant effect (Wilcoxon signed rank test).

We start by validating our method using synthetic behavior (for which we know the ground truth) and show that at test time, without further optimization, the network successfully recovers unseen agents’ RL parameters. We then present a proof-of-concept using two existing empirical datasets [9, 22], where human individuals performed a two-armed bandit task. For both datasets, we used t-RNN that was initially trained using synthetic data. Then, the network’s weights were fixed, and we applied it to predict individuals’ actions and to infer trial-by-trial theoretical RL parameters. In the first dataset, we utilized t-RNN to describe higher volatility in theoretical parameters for psychiatric individuals compared with healthy individuals, a group known for unstable internal mental states [23]. In the second dataset, we illustrated that t-RNN was able to capture dynamical changes in individuals’ directed and random exploration strategies [24]. Remarkably, across both datasets, we showcased that t-RNN produces parameter estimations that are meaningful while maintaining a level of predictive accuracy that is higher than theoretical models using a maximum-likelihood estimation approach [1]. Overall, we contend that our method can be used as a general framework, applicable to various theoretical models of behavior and datasets, and we discuss its advantages in enhancing the expressiveness of theoretical models to capture complex non-stationary behavior.

## Results

In the current study, we present a novel approach to simultaneously predict human behavioral choices and generate trial-by-trial estimations of theoretical RL parameters using neural networks. To demonstrate this approach, we use the well-known two-armed bandit task widely used in human neuroscience to investigate cognitive and neural processes. Initially, we train a t-RNN on artificial data obtained from theoretical RL agents performing the task and then keep the weights fixed to predict unseen behavior. We validate the effectiveness of our approach using synthetic data, followed by two use cases where we demonstrate t-RNN’s ability to predict actions and infer time-varying theoretical RL parameters in empirical datasets where human individuals performed the same task.

### Theoretical recurrent neural network (t-RNN) framework

Here we present a proposed method we termed t-RNN and describe how the network is trained to infer underlying theoretical RL parameters from behavioral observation of RL agents engaged in a two-armed bandit.

#### Two-armed bandit task

We present our model in the context of a generic two-armed bandit task widely used decision-making task to study behavior [1, 15]. At each trial of time *t*, the agent is forced to choose between two possible actions, *a* (denoted *L* and *R*). Next, a stochastic reward, *r*, is delivered. The reward probability of each action is fixed during a block of *B* trials and selected randomly at the beginning of each block from a pre-existing setting (see Materials and methods for more details).

#### t-RNN architecture

Our model as depicted in Fig. 1F, is based on a recurrent neural network (RNN) consisting of fully connected (FC) layers and gated recurrent units (GRU; [25]). At each step, the inputs to the t-RNN are the two observed time-dependent variables, the action *a*_*t*_ and the reward *r*_*t*_, and it produces two outputs: currents estimate of the agent RL parameters 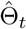 (Θ denotes a vector of RL parameters; subscripts *t* denotes time-varying RL parameters), and prediction of the future action 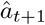.

#### t-RNN training

The two outputs that t-RNN produces, namely the hidden RL parameters and future action, are processed through different loss terms. The RL parameters are predicted in two stages. First as a multiclass classification task, after employing quantization of each RL parameter to a discretized space (i.e., ℝ *→*ℕ; see Materials and methods), and then as a regression problem. It is important to note that quantization was used only to stabilize the network training, while the network’s final output remains continuous. This procedure is based on previous work [26, 27] which has shown that MSE loss is much harder to optimize and often converges to a fixed output that reflects the mean. To assure that quantization did not affect our performance in any meaningful way, we ran multiple experiments in which we varied the number of the RL parameters quantization and found similar results (see S1 Table in the Supporting Information).

In the first stage, *a*_*t*_ (coded using one-hot transformation) and *r*_*t*_ is passed through a single layer of 32 GRU cells and the output is projected to *m* = |Θ| FC layers with ReLU activation to obtain the network intermediate prediction of the categorical class of each RL parameter. We then compute a cross-entropy loss (denoted as 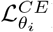) with regard to the true class for each RL parameter separately.

Next, to allow a continuous recovery of each RL parameter, in the second stage, the network categorical class prediction (i.e., the logits) is passed through *m* additional regression heads (each FC layer of size 1; each FC layer is responsible for a different RL parameter). We compute the mean-square error (denoted as 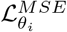) with regard to the true continuous values of each RL parameter separately.

In addition to estimating the parameters, our network also predicts the future action, *a*_*t*+1_. For this aim, we concatenate the network categorical class predictions (i.e., the logits) together with *a*_*t*_ and *r*_*t*_ and passed this vector through a second single layer of 32 GRU cells. We project the output to an FC layer with SoftMax activation to obtain a probability distribution over possible future actions and computed a cross-entropy loss (denoted as 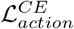) with regards to the true future action *a*_*t*+1_.

In the Supporting Information, we provide an additional experiment to examine the necessity of concatenation and its impact on the results. To investigate this, we employed ablation techniques and trained a version of the t-RNN model without concatenation (see Fig. S13 for further details).

We now define the combined loss function used to train the network,

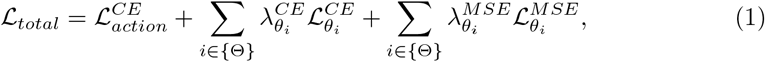

where *λ*^*CE*^ and *λ*^*MSE*^ are hyperparameters, set early during the development process such that each loss term is approximately of equal magnitude (see Materials and methods for further details).

### Validating t-RNN using synthetic behavior

We first demonstrate our method on simulated data of RL agents for which the ground truth RL parameters are known. Specifically, we trained t-RNN to predict the actions and parameters of a theoretical RL model. We then generated test agents on which the network was not trained with and compared the performance of t-RNN, to both stationary theoretical RL model (using maximum-likelihood [1]) and Bayesian particle filtering [28].

#### Theoretical RL model

To generate a training set of t-RNN, we simulated Q-learning [29, 30] agents performing a two-armed bandit task. Specifically, for the Q-learning agents, at each experience of time *t*, the agent updates the Q-values (initialized to zero at the beginning of each block) of the action selected *a*_*t*_ using the obtained reward *r*_*t*_ by,

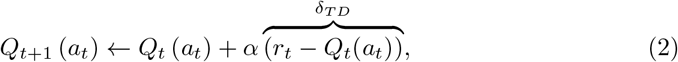

where 0 *≤ α ≤* 1 is a learning-rate free-parameter, which determines the extent to which newly acquired information overrides old information, and *δ*_*T D*_ is the temporal-difference (TD) error that drives learning.

These Q-values are in turn transformed to a stochastic choice-rule policy in accordance with the Boltzmann distribution,

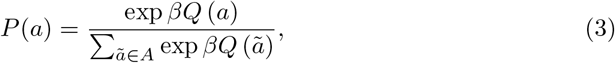

where *β ≥* 0 is the inverse-temperature free-parameter, which determines the randomness of action selection.

Training t-RNN was done with a synthetic training set of 2,000 simulated Q-learner agents operating with Eq. 2 and 3. Each agent performed a two-armed bandit task for 10 blocks of 100 trials each, and the agents’ RL parameters were sampled from a uniform distribution. Importantly, to stabilize the training procedure, we applied two techniques. First, to allow for continuous recovery of each RL parameter, we performed a quantization preprocessing step, where we discretized each RL parameter into evenly spaced bins, which acted as an intermediate prediction target during training. Second, to facilitate t-RNN’s flexibility to recover time-varying RL parameters, we simulated agents with non-stationary parameters. For each agent, there was a small probability per trial that the agent’s parameters would be re-sampled from the same uniform distribution (see Materials and methods for more details).

For comparison purposes, we included two additional baseline methods for estimating hidden RL parameters. The first baseline method was a stationary Q-learning model (denoted Q-stationary), in which stationary RL parameters were estimated for each agent individually, using a maximum-likelihood estimation. The second baseline method was Bayesian particle filtering (denoted Bayesian) adopted from [28]. Here, underlying RL parameters are assumed to drift slowly according to a Gaussian random walk and are estimated for each agent on a trial-by-trial basis using Bayesian particle filtering (see Materials and methods for more details).

#### Results

To validate our method, after training t-RNN, we simulated 30 additional test agents (Materials and methods) on which the network was not trained and recorded the predictions made by t-RNN as well as the two baseline methods. First, we calculated the binary cross-entropy (BCE) between the true actions and predicted actions of each of the three models (stationary Q-learning, Bayesian, and t-RNN). We found that t-RNN had the lowest error compared to the two other models (*p <* .01 when comparing Q-stationary and t-RNN using Wilcoxon signed rank test, and *p <* .05 when comparing Bayesian and t-RNN; see BCE_*action*_ in Fig. 2A;). Second, we calculated the mean-squared error (MSE) between the true and estimated RL parameters and found that t-RNN outperformed the two alternatives methods in terms of MSE_*β*_ (*p <* .01 when comparing Q-stationary and Bayesian to t-RNN using Wilcoxon signed rank test; see Fig. 2A). Note that although t-RNN exhibited the lowest MSE*α* compared to both baseline methods, it was not significant. This finding confirmed that our network can accurately recover hidden RL parameters of test agents operating with various underlying RL parameter dynamics (see S2 Table for raw results).

**Fig 2.**
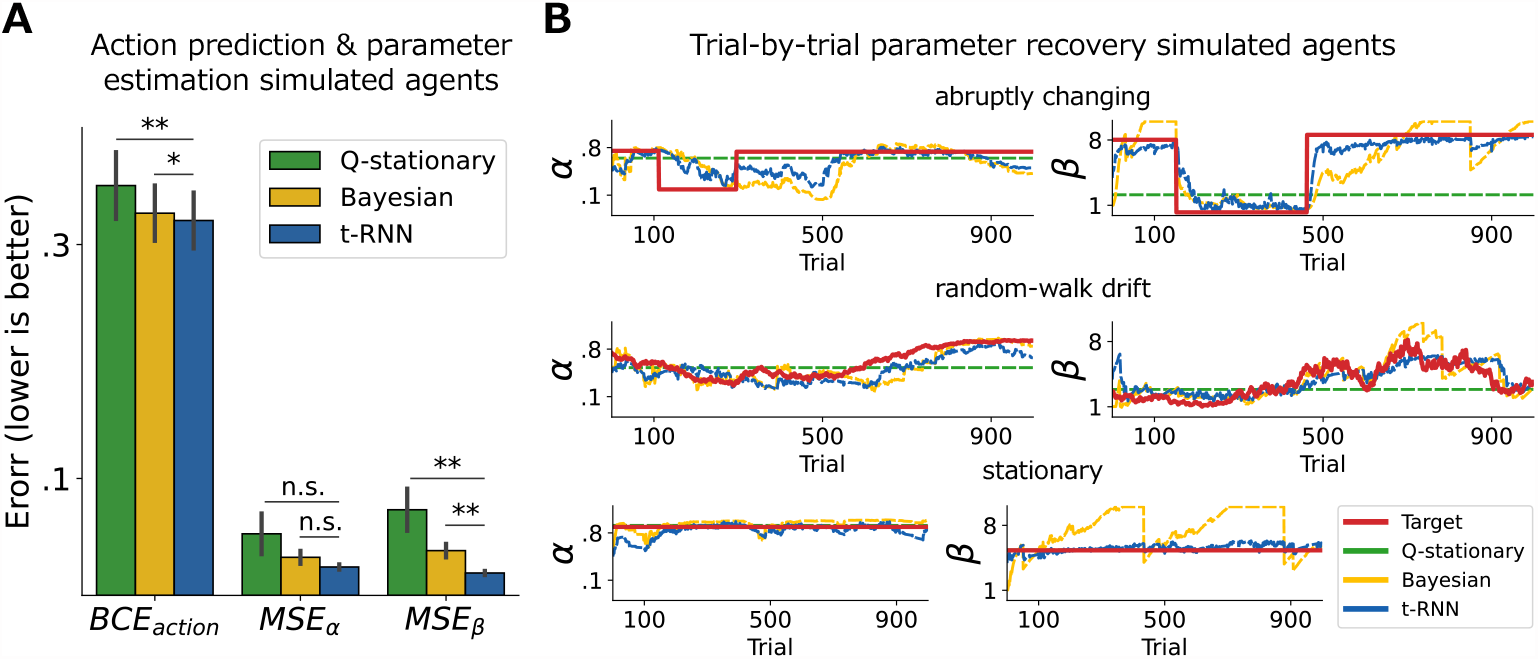
Validating the ability of t-RNN to predict both agents’ actions and to infer RL parameters of simulated data. **(A)** Action prediction (measured with binary cross entropy; BCE) and RL parameter estimation error (measured with mean-square error; MSE) averaged across 30 artificial test agents (black lines indicate s.e.m). We found that our method exceeded alternative methods (Q-stationary [Maximum-likelihood], green; Bayesian [particle filtering], yellow) in both action prediction (left) and parameter estimation (middle and right). **(B)** Recovery of theoretical RL *α* learning-rate (left) and *β* inverse-temperature (right) parameters of three example simulated agents with different RL parameter trajectories. These trajectories included abruptly changing (top), random-walk drift (middle), and stationary (bottom) RL parameters. Overall, we found that our network (blue dotted line) was able to effectively recover the true dynamic RL parameters (red solid line). This demonstrates the versatile ability of t-RNN to accurately capture a wide range of time-varying RL parameter dynamics. Significant effect at * *p <* .05, ** *p <* .01; n.s., a nonsignificant effect (Wilcoxon signed rank test).

To further examine and illustrate the t-RNN capabilities, we plotted (see Fig. 2B) the true and recovered RL parameters of three types of test agents: abruptly changing RL parameters (top), gradually changing RL parameters (middle), and agents with stationary RL parameters (bottom). Visual inspection of Fig. 2B indicates that t-RNN generated predictions that closely matched the true RL parameter trajectories. Overall, these results indicate t-RNN flexible ability to recover underlying RL parameters while keeping high predictive accuracy. In the Supporting Information, we present two additional experiments. The first experiment explores the recovery of agents simulated in a volatile environment (see S3 Table), where we observed that t-RNN maintains superior performance compared to both baseline methods. In the second experiment, we examine the ability of t-RNN to estimate stable parameters compared to the stationary Q-learning method as a function of the number of observations (see S4). Our findings demonstrate that t-RNN performs well for stable parameters and is comparable to the stationary method when more than one parameter is included in the theoretical model. Moreover, for a low number of observations (below 50 trials), t-RNN outperforms the stationary estimation.

### Using t-RNN to interpret behavior of psychiatric individuals

After confirming the ability of our method to recover hidden RL parameters with a variety of test agents, we next present an application of t-RNN to an empirical behavioral dataset collected by Dezfouli et al., 2019 [9] where 33 individuals diagnosed with bipolar disorder, 34 with depression and 34 individuals who were matched healthy controls (total of *N* = 101) performed two-armed bandit task (see Materials and methods for more details).

#### Theoretical RL model

To model subjects’ behavior (and to train t-RNN), we consider the same RL Q-learning model as mentioned above (see Eq. 2 and 3) with an additional action preservation free-parameter added to the SoftMax choice-rule policy,

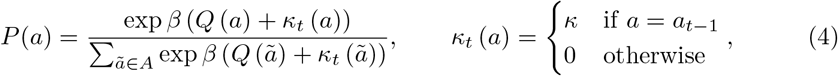

where *−*0.5 *≤ κ ≤* 0.5 is an action preservation free-parameter determining the agent’s tendency to stick with (or switch) a previous action regardless of the reward [31].

#### Results

Following training of the t-RNN with a synthetic training set of Q-learner agents using Eq. 2 and 4, we fixed the network’s weights and recorded the predictions made by the t-RNN on the behavioral dataset of humans who performed a two-armed bandit task [9]. Additionally, we fitted a stationary Q-learning with preservation model (denoted QP-stationary) to each subject, which estimated stationary RL parameters via maximum-likelihood estimation, and a Bayesian model fitted with particle filtering. Furthermore, we fitted a data-driven RNN (d-RNN) to subject behavior, as described in [9], which solely produced action predictions (see Materials and methods for more details).

As expected, we found that t-RNN was able to maintain a higher action prediction of human behavior compared to the QP-stationary model but not compared to d-RNN (see Fig. 1B; *p <* .05 when comparing QP-stationary and t-RNN using Wilcoxon signed rank test, and *p <* .0005 when comparing t-RNN and d-RNN). To further explore this result, we divided the subjects according to their diagnostic groups and examined the action predictions within each group. We found that t-RNN superiority over QP-stationary and Bayesian models was mainly due to the improved action prediction of bipolar and depressed individuals (*p <* .01 comparing QP-stationary and Bayesian to t-RNN in the bipolar group; *p <* .01 comparing QP-stationary to t-RNN and *p <* .001 comparing Bayesian to t-RNN in the depressed group; Wilcoxon signed rank test), while there was no significant difference in its prediction accuracy for healthy controls (*p* = 0.36 comparing QP-stationary to t-RNN; *p* = 0.07 comparing Bayesian to t-RNN; see Fig. 3A and S5 Table for the raw results).

**Fig 3.**
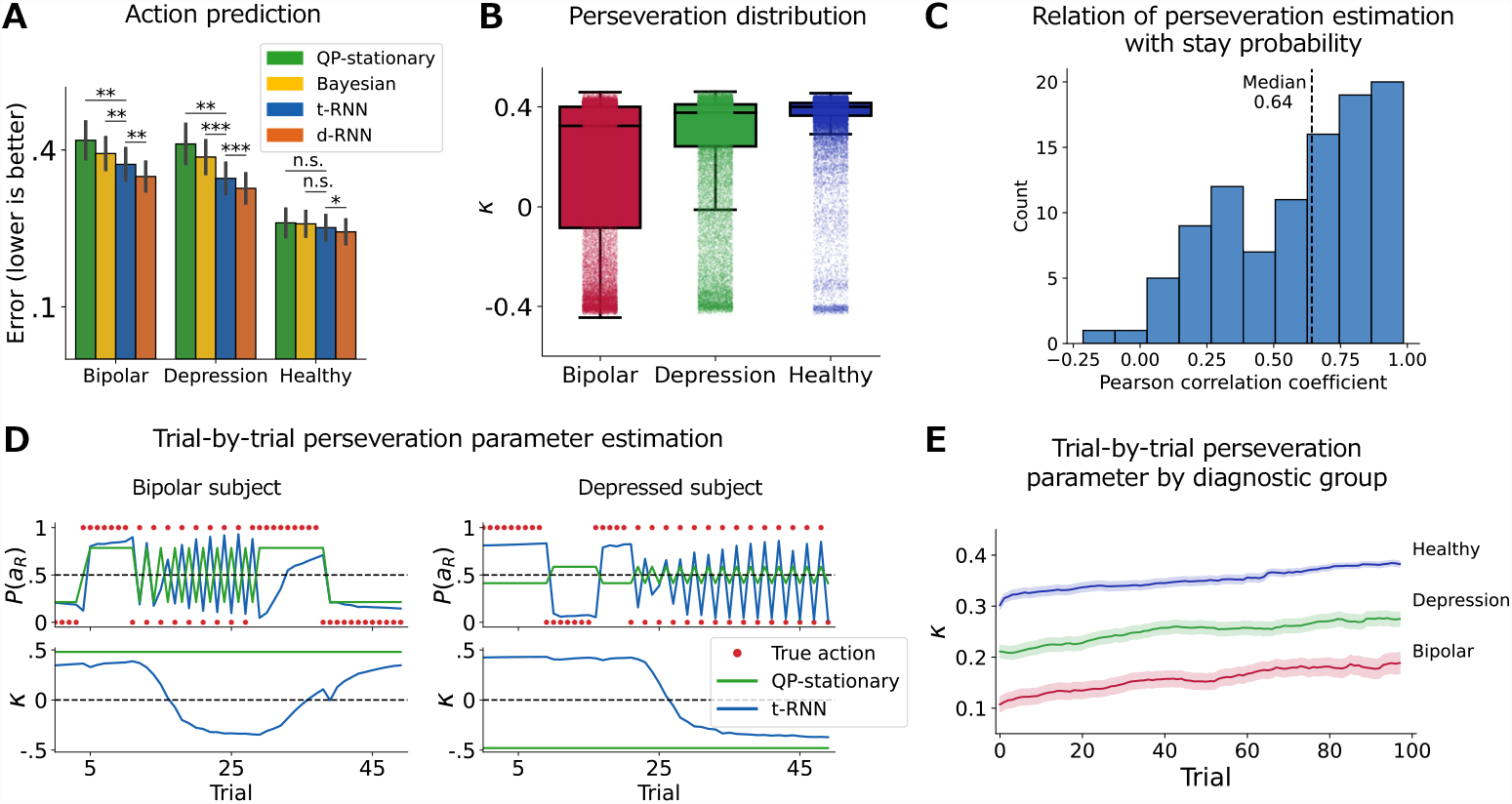
Application of t-RNN to behavioral data of psychiatric and non-psychiatric individuals [9]. **(A)** Action prediction (measured with BCE; black lines indicate s.e.m) divided by diagnostic labels. Bipolar and depressed subjects at the group level are explained significantly better by t-RNN (blue) compared to the QP-stationary model (green) which assumes that the RL parameters are fixed, as well as to the Bayesian model (yellow). **(B)** Boxplot of the time-varying *κ* preservation parameter estimates by t-RNN across the three diagnostic groups (middle black solid lines denote the median; light dots indicate a single trial estimate of the *κ* preservation parameter). Result suggests higher volatility in the clinical groups mainly the bipolar group, compared with the healthy group. **(C)** Distribution of the Pearson correlation between stay probability (calculated using moving average of 10 trials) and time-varying *κ* preservation parameter produced by t-RNN for each subject individually. Result shows a strong relation between t-RNN time-varying RL *κ* parameters estimation and moving average of the stay probability. **(D)** Sequence of selected actions of two example subjects, with bipolar on the left and depressed on the right. The top panel shows the action prediction, and the bottom panel shows the RL *κ* parameter estimation produced by t-RNN (blue) and QP-stationary model (green) for a 50-trial segment (see Fig S10 for all trials). Both subjects seemed to switch between selecting the same action for several trials to repeatedly alternating between both actions every trial (red dots). We found that the t-RNN model was able to detect this visually apparent shift in behavior, estimating a change in the *κ* action perseveration parameter. In contrast, the QP-stationary model was not able to capture this transition. **(E)** Trial-by-trial *κ* preservation parameter estimates by t-RNN averaged over the first 100 trials of each block and for each diagnostic group separately (shaded area signifies the s.e.m). In all three groups, subjects show a steady increase in their tendency to perseverate their actions as the block progresses. Significant effect at * *p <* .05, ** *p <* .01, *** *p <* .001; n.s., a nonsignificant effect (Wilcoxon signed rank test).

We next turn to explore the dynamic estimation of the theoretical parameters. First, we assessed the volatility in *κ* preservation estimates estimated trial-by-trial using t-RNN. Our results revealed that the bipolar group exhibited lower values and greater volatility in these estimates compared to the healthy group (*M* = 0.15, *SD* = 0.30 for the bipolar group, *M* = 0.35, *SD* = 0.14 for the healthy group; see Fig. 3B). This finding might correspond with the clinical nature of the disorder where volatility is a main feature [23]. However, for the aim of the current study, this is mainly a proof-of-concept to the t-RNN’s ability to inform us regarding volatility in underlying parameters, while also providing better action predictions compared with a stationary QP-stationary model. In S6 Table, we show additional distribution of all RL parameters.

Second, to gain insight into the association between RL parameter estimation and observed behavior, we selected two clinical participants with bipolar disorder and depression whose behavior displayed clear transitions between high (sticking with the same action) and low (switching the action selection every trial) levels of perseveration. We plotted the sequence of selected actions alongside the action predictions and inferred *κ* estimates generated by t-RNN, as well as the predictions of the QP-stationary model for a 50-trial segment (see Fig. 3D and Fig. S10 for all trials). We found that the observed change in action perseveration was closely coupled with a change in the *κ* parameter, and the t-RNN model accurately detected the visually apparent shift in behavior. In contrast, the stationary model, which assumes fixed RL parameters, was unable to capture this transition, resulting in lower action prediction accuracy. These results highlight the utility of t-RNN in capturing dynamic changes in RL parameters and their association with observed behavior.

Third, to further support that t-RNN parameters estimation indeed corresponds well with changes in observed behavior across all individuals, we turn to a well-established and easy-to-interpret model-agnostic measurement and calculated a moving average of the stay probabilities (probability of repeating the previous action; window size of 10 trials). If indeed the t-RNN *κ* perseverate dynamic parameter is doing a good job of capturing fluctuations in perseveration, we should expect a high correlation with the moving average model-agnostic estimate. We calculated a Pearson correlation between the moving average and t-RNN *κ* perseverate trial-by-trial estimation for each individual (see Fig. 3C). Overall, we found that the average Pearson across individuals was very high (Median= 0.64; see S7. Fig for further detailed examples). In the Supporting Information we provide a similar analysis with the *β* inverse-temperature parameter (see S8. Fig)

Finally, we plotted the trial-by-trial *κ* perseveration average for each group, showing that overall, across groups individuals increased their tendency to perseverate with the previous action selection which might reflect an overall reduction motivation as the block progresses (see Fig. 3E). Overall, we were able to show that t-RNN is able to produce trial-by-trial estimates of RL parameters, while also keeping higher predictive accuracy compared with QP-stationary model. In the current dataset, we provided a proof-of-concept on how t-RNN might be able to inform of higher volatility in RL parameters in clinical psychiatric groups.

### Using t-RNN to interpret exploration behavior of human individuals

To demonstrate the generality of our proposed method, we present a second application of t-RNN to empirical behavioral dataset collected by Gershman, 2018 [22] where *N* = 44 humans performed two-armed bandit task to examine human exploration strategies.

#### Theoretical RL model

To model subjects’ behavior, we consider the same model hybrid exploration model presented in Gershman, 2018 [22]. The learning model included a Kalman filtering algorithm [15, 32] where agents recursively compute the posterior mean and variance for each action by,

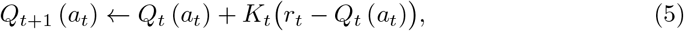

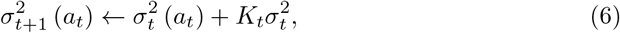

where *Q*_*t*+1_ (*a*) and 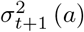 is the posterior mean and variance estimates over the value of action *a*, respectively and *K*_*t*_ is the Kalman gain given by,

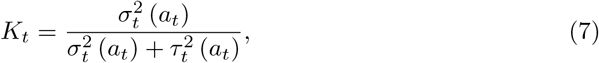

where 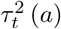 is the error variance. At the beginning of each block the initial values estimate for both actions were set to the prior mean, namely, *Q*_0_ (*a*) = 0, 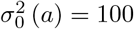 and 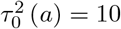 (see Materials and methods for more details).

Action values estimate were transformed to action policy with a hybrid of random and directed exploration [24] choice-rule,

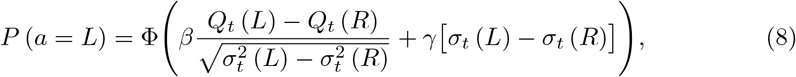

where Φ is the cumulative distribution function (CDF) of the standard Gaussian distribution (i.e., probit), 0 *≤ β ≤* 4 is a free-parameter that controls the contributions of random exploration (similar in spirit to the inverse-temperature parameter of the SoftMax choice-rule of Eq. 3) and 0 *≤ γ ≤* 1 is a free-parameter that controls the contributions of directed exploration.

This model can also be interpreted as a combination of Thompson sampling [33] and UCB policies [34], where the first term in the brackets promotes a random exploration that is proportional to the difference in the value estimates, *Q*_*t*_ (*L*) *− Q*_*t*_ (*R*) together with the total uncertainty 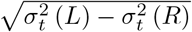 much like Thompson sampling. The second term in the brackets, *σ*_*t*_ (*L*) *− σ*_*t*_ (*R*), is the so-called relative uncertainty, synonyms with UCB policy which promotes a directed exploration toward the less certain option (i.e., information bonus).

#### Results

After the training of the t-RNN with a synthetic training set of Q-learner agents using Eq. 5, 6, 7 and 8, we fixed the network’s weights and recorded the predictions made by the t-RNN on the behavioral dataset of humans who performed a two-armed bandit task [22]. Additionally, we fitted a stationary hybrid exploration model, a Bayesian model, and a d-RNN to subject behavior (Materials and methods). We found significantly higher action prediction for t-RNN compared to both the stationary hybrid exploration model (*p <* .05; Wilcoxon signed rank test) and the Bayesian model (*p <* .01), but no significant difference in prediction accuracy when comparing t-RNN to d-RNN (*p* = 0.12; see Fig. 4A and S11 Table for raw results).

**Fig 4.**
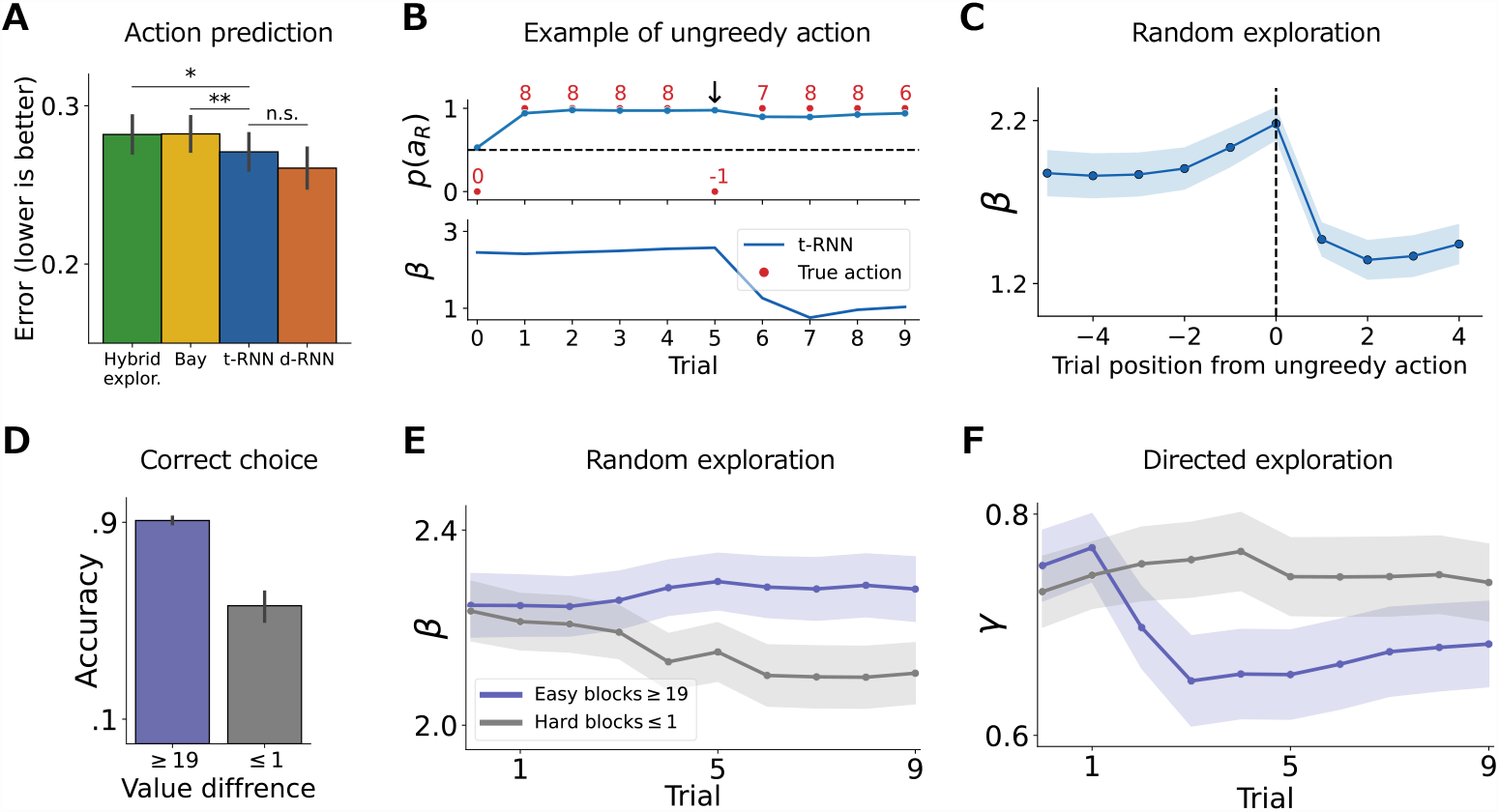
Application of t-RNN to investigate changes in random and directed exploration [22]. **(A)** Action prediction (measured with binary cross entropy; BCE; black lines indicate s.e.m). We found a significant improvement in the action prediction of the t-RNN model (blue) compared to the hybrid exploration model (green; * *p <* .05) and the Bayesian (yellow; ** *p <* .01). **(B)** Example of a single instantiation of ungreedy action. The top panel depicts the sequence of chosen actions, obtain rewards, and t-RNN action predictions as red dots, red numbers, and blue solid lines, respectively. The subject consecutively chooses the higher value action and suddenly switches and chooses the lower value action (denoted with a downward arrow). The bottom panel depicts t-RNN *β* random exploration parameter estimation, where it estimates a step decrease in the parameter after the ungreedy action selection. **(C)** Dynamics of the random exploration parameter averaged across all instantiations of ungreedy action. t-RNN estimates a sharp and sudden decrease in the random exploration parameter after an ungreedy action is taken. **(D)** Correct choice rate of individuals’ actions divided by blocks where the true value difference is high (*≥* 19; easy blocks; purple) and blocks where the true value difference is low (*≤* 1; hard blocks; grey). Subjects are more accurate when it is easy to distinguish the correct action compared to when it is hard. **(E)** Dynamics of the *β* random exploration parameter estimation produced by t-RNN average overall easy (purple) and hard (grey) blocks. Random exploration increases in hard blocks (grey) where it is difficult to tell apart which action is best, whereas in the easy blocks (purple) random exploration decreases as subjects become more deterministic in their action selection. **(F)** Dynamics of the *γ* directed exploration parameter average overall easy (purple) and hard (grey) blocks. Directed exploration is high at the beginning of the block where uncertainty is equal in both conditions, however only in the easy blocks (purple) where it is easy to differentiate between the better/worst arm, directed exploration decreases as the block evolves. In panel C, E and F solid line/dots signifies the mean estimation and the shaded area correspond to s.e.m.

To investigate the trial-by-trial dynamics of the t-RNN’s theoretical parameter estimation, we conducted a manual search of the data to identify an instance in which an individual made an ungreedy action. Specifically, we located a trial where an individual chose option L despite having been rewarded for option R for the previous four consecutive trials. We observed a sharp decrease in the t-RNN’s *β* random exploration parameter estimation, indicating its sensitivity to changes in observed behavior (see Fig. 4B). To determine whether this pattern generalizes across individuals and trials, we aggregated all trials (a total of 32 trials) in which individuals selected the better rewarding arm for three or more consecutive trials, were then rewarded, and subsequently switched to the alternative, poorer arm. Our analysis revealed an overall decrease in the *β* random exploration parameter produced by t-RNN in these cases (mean *±* s.e.m. before: 2.1 *±* 0.079, after: 1.40 *±* 0.080; see Fig. 4C). In the Supporting Information we provide a similar example for the *γ* directed exploration parameter (see S12. Fig).

Next, we investigated the overall dynamics of the two exploration parameters (random and directed) within blocks. Specifically, we examined blocks where the true expected value difference between the two arms was above or equal to 19 (referred to as “easy” blocks; total of 91 blocks) and blocks where the true expected value difference between the two arms was below or equal to 1 (referred to as “hard” blocks; total of 110 blocks). It is worth noting that participants were not informed whether a block was easy or hard and were required to start with a similar strategy in the first trial where the difficulty of the block was unknown. We hypothesized that participants would change their exploration strategy across time, and predicted that in easy blocks, individuals would reduce random exploration since the best action is easy to detect, whereas, in hard blocks, individuals would show a higher tendency for random exploration (indicating either noisy behavior due to block difficulty or a desire to test the arms in a more heterogeneous fashion to gain more information, where the information gain is relatively small for the participant).

We found that participants had a higher correct choice rate in easy blocks compared to hard blocks (mea *±* s.e.m. easy: 0.90 *±* 0.007, hard: 0.56 *±* 0.003; see Fig. 4D). Consistent with our hypothesis, t-RNN estimated an increase in random exploration (i.e., a decrease in the *β* parameter) across hard blocks and a decrease across easy blocks (see Fig. 4E). We also observed the same pattern for the *γ* directed exploration, where overall directed exploration was high at the beginning of the block, possibly reflecting the individual’s lack of knowledge about the environment. However, there was a selective decrease in directed exploration only for easy blocks, where the best action was easy to locate (see Fig. 4F).

Our results indicate that individuals modify their exploration strategies based on both the block difficulty and phase, as evidenced by changes in both random and directed exploration parameters. Importantly, t-RNN was successful in capturing these nuanced changes in exploration strategies, thus providing a second proof of the utility of our method.

## Discussion

In this work, we introduce the notion of a “theoretical-RNN” (i.e., t-RNN), where neural networks are utilized to augment the expressiveness of theoretical models of behavior. Specifically, we train an RNN on artificial data simulated from a theoretical model, freeze the network weights, and use the network to predict trial-by-trial theoretical RL parameters and future actions. The immediate benefit of this approach is its ability to infer non-stationary trial-by-trial theoretical parameters, thus allowing us to examine dynamics in latent parameters thought to underlie human behavior. We first validated our framework with simulated data, showing our ability to recover both stationary and non-stationary theoretical parameters. We then applied this approach to two independent empirical datasets. In the first dataset, we demonstrated that t-RNN can capture the volatile behavior patterns of psychiatric patients, which differ significantly from healthy controls. In the second dataset, we show that t-RNN can identify the changes in human exploration strategies that are modulated by task difficulty and block phase.

Several studies have explored the benefits of using neural networks fitted directly to behavioral data to examine various behavioral phenomena, including response time in task switching paradigms [35], the identification of novel theories in behavioral economics under risky choice [19], and particularly relevant to the current research, reinforcement learning (RL) tasks [9, 18, 21, 36]. For example, Dezfouli et al., 2019 [9] trained a recurrent neural network (RNN) to predict future human actions in a two-armed bandit task which was found to outperform a standard RL model in pure action prediction. This result was also generalized to other, more complicated decision-making tasks, such as the four-armed bandit [36] and build-your-own-stimulus tasks [18]. Additionally, in [37, 38] the authors employed a feedforward neural network fed with inputs such as mouse stress, affective phenotype, and prior task performance to examine how neurological measures control RL parameters. More recently, in order to study individuals’ differences, [21] developed an autoencoder model, where an RNN encoder was used to map each agent’s behavior into a low-dimensional latent space. The low-dimensional latent space produces a disentangled representation that can be used to study variations in participants’ decision-making strategies. Overall, previous work shows the high ability of neural networks to learn to some extent the latent data generating model of human behavior across different paradigms. However, the main challenge remains the low interpretability of these methods.

Here, we consider our research as an expansion of the growing trend in using neural network models to study behavior and identify four distinct contributions of using a t-RNN framework. First, our method enhances the utilization of existing theoretical RL models, leveraging parameters with clear and straightforward interpretations. This allows us to dynamically estimate latent RL parameters based on empirical choice behavior. Additionally, we demonstrate that this approach exceeds stationary theoretical models fitted with conventional approaches (e.g., maximum-likelihood) by providing better predictive accuracy. Second, a known difficulty when training neural networks with empirical data is the need to obtain a large training set. Here, we demonstrate the validity of training RNN using synthetic data which can be arbitrarily large. Post training, and without further optimization, the trained network is then used to infer empirical behavioral data. Third, our method also holds the benefit that it can be applied directly and online to newly acquired data, thus providing solutions for latent parameter estimation in real time. This allows for online monitoring of behavior during cognitive tasks, facilitating personalized interventions or manipulations based on each individual’s current state. Fourth, the generality of our framework may be extended beyond choice behavior by utilizing theoretical models of other modalities. For example, given reaction times observation, we might be able to employ t-RNN to predict the dynamic theoretical parameters of evidence accumulation models. Another possibility is to utilize theoretical models of neural data (e.g., attractor networks) to train t-RNN in predicting latent theoretical parameters from the neural recordings.

An important benefit of t-RNN is its ability to predict theoretical parameters in a dynamic manner. While some recent studies have recognized the importance of relaxing the assumption of stationarity parameters in behavioral data [39–42], it remains common to treat behavior as a stationary process. Specifically, most of the computational modeling work in behavioral RL studies is performed by fitting a single set of parameters to each participant. Here, we compare our t-RNN framework to the Bayesian particle filtering method used in [14, 28, 43] which estimates hidden RL parameters assumed to slowly drift according to a Gaussian random walk. Our approach requires less prior specification (i.e., manual specification of prior and hyper-prior distributions) and provides a more flexible way to recover a wider range of RL theoretical parameter trajectories.

Another important aspect in the current work is the fact that a specific generative theoretical model, which serves as the supervised training signal for t-RNN needs to be specified. If human behavior is governed by a substantially different model, our method may have difficulties in accurately predicting behavior. Therefore, t-RNN is limited in its ability to predict parameters and actions, to the extent that the RL theoretical model it was introduced to during training accurately describes the participant’s behavior. However, the high flexibility of neural networks opens the way for using t-RNN in much more sophisticated ways in the future. For example, future studies can train t-RNN on simulated data that were obtained from different agents acting according to different models. Moreover, these agents might switch from one model to the other arbitrarily. This might allow the dynamic trial-by-trial estimation of both parameters and models that might generate the observed behavior, and allow to estimate moments where the subject is switching from one to another model. Furthermore, since t-RNN is trained on synthetic data that can be infinitely large, it might be possible to introduce during training a vast model space and then predict the most suitable model, parameter, and action dynamically for a single empirical dataset. Thus, by incorporating all three predictions into the training process (models, parameters, and actions), future studies might be able to better address the issue of model misspecification, as a single t-RNN model may have the ability to predict various theoretical models.

Some limitations of our work. First, while our study demonstrated the ability of t-RNN to capture the high temporal dynamics of theoretical parameters, we did not provide a detailed explanation for the underlying nature of such changes. Although our current study did not include biological measurements, we believe that our approach might be able to facilitate future research where changes in latent parameters can be linked to meaningful biological indicators. Integrating the t-RNN methodology with these indicators may offer a deeper understanding of the neural correlates of behavior and the interplay between latent parameters and physiological processes. Second, we acknowledge that the training settings can influence t-RNN performance, our study aims to present a proof-of-concept for the primary framework of using RNNs to predict theoretical parameters, with further work required to test the impact of different training settings on t-RNN performance. Lastly, it is interesting to speculate how the complexity of the theoretical model might affect our methods. Previous studies have demonstrated that increasing model complexity may be necessary to effectively capture a broader range of animal and human behaviors [44]. We hypothesize that for more complex theoretical models we will find an advantage for t-RNN, as neural networks are known to be able to learn complex non-linear mappings [45]. Additionally, it is plausible that under certain conditions, t-RNN may estimate parameters with fewer observations compared to more traditional approaches (see Fig. S4). However, further studies are required to ensure that our approach generalizes across a wide range of model complexities.

In conclusion, we argue that the advantages of our method outweigh its limitations and that we present a novel and versatile framework for enhancing the expressiveness of theoretical models to capture complex non-stationary behavior using neural networks. We believe that t-RNN serves as a bridge between traditional RL models and modern neural network models, allowing for a unified approach that can retain the strengths of both modeling paradigms.

## Materials and methods

### Code availability

The code is publicly available via: https://github.com/yoavger/harnessing_the_ flexibility_of_nn_to_predict_dynamical

### t-RNN Implementation

We implemented our network in PyTorch library [46], training with a single NVIDIA GeForce RTX 3080. We splitted the training set to 80/20, where 80% was used for training and the remaining 20% for validation. We use Adam optimizer [47] and a constant learning rate of 0.001. The number of hidden cells of the GRU layer was determined by a grid search selecting the one with the best validation loss which turn out to be 32 cells. We use early stopping together with dropout (*p* = 0.2 in both GRU layers) to prevent overfitting. We found that the best performance is obtained after a short training of fewer than 50 epochs. We also tested our model with different batch sizes (100,500,1000) and found the same results; therefore, we chose a batch sizes of 1000 to speed up the training time.

### Application of t-RNN to simulated data

#### Task structure

Agents performed a two-armed bandit task for 10 blocks of 100 trials each. At each trial the agent received a stochastic binary reward of *r* = 1 for rewarded trials, or *r* = 0 otherwise. The reward probability of each action is fixed during a block of 100 trials and is selected randomly at the beginning of each block from three possible Bernoulli distribution: 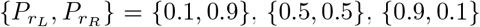.

#### t-RNN training

Training t-RNN was done with a synthetic training set of 2,000 simulated Q-learners agents operating with Eq. 2 and 3. Each agent performed a two-armed bandit task for 10 blocks of 100 trials each. Each agents RL parameters were sampled from a uniform distribution of *α ∼ U* (0, 1) and *β ∼ U* (0, 10). In the quantization preprocessing step, we discretized the *α* parameter into five evenly spaced bins (of size 0.2), and the *β* parameter into five evenly spaced bins (of size 2).

Importantly, to facilitate t-RNN flexibility to recover time-varying RL parameters, we simulated agents with non-stationary parameters. For each agent, there was a small probability of (*p* = 0.005) per trial that the agent’s parameters will be re-sampled from the same uniform distribution. To keep the set from changing too rapidly, the overall number of trials in which the parameters were re-sampled was limited to four, and after each trial of parameters re-sampling the probability of another change was fixed to zero for 100 trials.

The combined loss function (Eq. 1) included the following *λ* hyperparameters values: 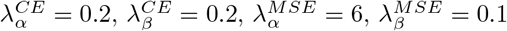, so that each loss term is approximately of equal magnitude.

#### Stationary Q-learning

Here, RL parameters were estimated for each agent individually, using a maximum-likelihood estimation approach (Python SciPy [48] minimized function with L-BFGS-B optimization, and five different starting search locations to avoid local maxima). This approach allowed for an estimation of a single and fixed set of RL parameters for each agent [1].

#### Bayesian particle filtering

Here, underlying RL parameters are assumed to drift slowly according to a Gaussian random walk,

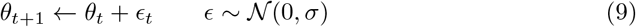

and are estimated for each agent on a trial-by-trial basis using Bayesian particle filtering. We implemented the same model as described by Samejima et al., 2003 [28]. To recover the simulated data, we set 1,000 particles to approximate the distribution of two hidden RL parameters {*α, β*}. Initial distribution of the particles were set to be Gaussians, with means of {0, 0} and variances of {3, 1}, respectively. We set the hyper-prior for the dynamics of *α* and *β* as *σ*_*α*_ = 0.1 and *σ*_*β*_ = 0.05, respectively (we also tested using hyper-prior as low as *σ*_*α*_*/σ*_*β*_ = 0.005*/*0.0005 and us high as *σ*_*α*_*/σ*_*β*_ = 0.05*/*0.005 and *σ*_*α*_*/σ*_*β*_ = 0.1*/*0.5, these setting resulted in poorer results). We also clipped the *β* to range between (0,10) to match the estimation range of other models.

#### Test agent

In order to validate and compare the performance of t-RNN against baseline methods, we simulated a test set of 30 artificial Q-learning agents (Eq. 2 and 3). Each agent was simulated on the same task structure, consisting of 10 blocks of 100 trials, and his RL parameters were sampled from a uniform distribution of *α∼ U* (0, 1) and *β ∼ U* (0, 10). The test set included three groups of 10 agents each. The first group had stationary RL parameters, while the second group had RL parameters that abruptly changed with a small probability for each trial (*p* = 0.005). The third group had RL parameters that drifted slowly according to a Gaussian random walk (Eq. 9) with sigma values of *σ*_*α*_ = 0.1 and *σ*_*β*_ = 0.05.

### Application of t-RNN to psychiatric human individuals

#### Behavioral dataset and task structure [9]

The behavioral dataset includes *N* = 101 individuals that performed the bandit task for 12 blocks, with a free-choice rate of 40 second per block. At each trial subjects received a stochastic binary reward of *r* = 1 for rewarded trials, or *r* = 0 otherwise. The reward probability of each action was fixed during a block and selected at the beginning of each block from six possible settings: 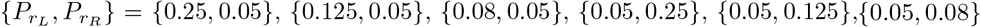. Each possible reward setting was repeated twice, for a total of 12 blocks. The data included task trials of 33 individuals diagnosed with bipolar disorder, 34 with depression and 34 individuals who were matched healthy controls (see Dezfouli et al., 2019 [9] for further details).

#### t-RNN training

Training t-RNN was done with a synthetic training set of 2,000 simulated Q-learners agents operating with Eq. 2 and 4. Each agent performed a two-armed bandit task for 12 blocks of 100 trials each. Each agents RL parameters were sampled from a uniform distribution of *α ∼ U* (0, 0.2), *β ∼ U* (0, 10) and *κ ∼ U* (− 0.5, 0.5). In the quantization preprocessing step, we discretized the *α* parameter into three evenly spaced bins (of size 0.066), the *β* parameter into five evenly spaced bins (of size 2) and the *κ* parameter into five evenly spaced bins (of size 0.2). Again, to facilitate t-RNN flexibility to recover time-varying RL parameters, we simulated agents with non-stationary parameters, with the same setting as described above.

The combined loss function (Eq. 1) included the following *λ* hyperparameters values: 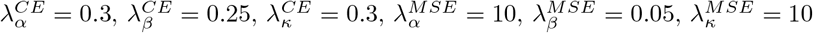, so that each loss term is approximately of equal magnitude.

#### d-RNN model

Data driven RNN was implemented following the same RNN model as described by Dezfouli et al., 2019 [9] (term here d-RNN). d-RNN is composed of a single GRU layer of 10 hidden units. The inputs to the d-RNN are the current action *a*_*t*_ and the reward *r*_*t*_ received after taking that action. The outputs are probabilities over the action of the next step *a*_*t*+1_. We train the d-RNN with and Adam optimizer, learning rate of 0.001 and batch size of 1000. We assess performance with the same leave-one-out cross-validation schema depicted in [9]. At each CV round, one of the subjects was withheld and d-RNN was trained using data from the remaining subjects. Importantly, this procedure is done separately for each diagnostic group. For example, in order to predict the actions of a withheld bipolar subject, we use data from only the remaining bipolar subjects to train the d-RNN model.

#### Stationary Q-learning with preservation (QP-stationary)

Here, stationary RL parameters are estimated for each subject individually, with maximum-likelihood estimation (Python SciPy [48] minimized function with L-BFGS-B optimization, and five different starting search locations to avoid local maxima).

#### Bayesian particle filtering

For each individual, we set 1000 particles to approximate the distribution of three hidden RL parameters {*α, β, κ*}, Gaussian with means of {*−*2, 1, 0} and variances of {1.5, 1, 1.5}, respectively. The hyper-prior for the dynamics of *σ*_*α*_ = 0.05, *σ*_*β*_ = 0.005, and *σ*_*κ*_ = 0.05. We clipped the beta to range between (0,10) to match the estimation range of other models.

### Application of t-RNN to interpret exploration behavior of human individuals

#### Behavioral dataset and task structure [22]

The behavioral dataset includes *N* = 44 individuals that performed the bandit task for 20 blocks of 10 trials each. At each trial subjects received a stochastic reward drawn from a Gaussian distribution with means of *μ*_*L*_,*μ*_*R*_ (for actions *L* and *R* respectively) and a variance (i.e., error variance) of 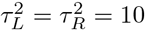. The mean reward probability of each action was fixed during a block and was drawn at the beginning of each block from a Gaussian distribution, with mean 0 and variance 100 (see Gershman, 2018 [22] for further details).

#### t-RNN training

Training t-RNN was done with a synthetic training set of 10,000 simulated Q-learners agents operating with Eq. 5, 6, 7 and 8. Each agent performed a two-armed bandit task for 20 blocks of 10 trials each. Each agents RL parameters were sampled from a uniform distribution of *β∼ U* (0, 4) and *γ ∼ U* (0, 1). In the quantization preprocessing step, we discretized the *β* parameter into five evenly spaced bins (of size 0.8), the *γ* parameter into five evenly spaced bins (of size 0.2). Again, to facilitate t-RNN flexibility to recover time-varying RL parameters, we simulated agents with non-stationary parameters, with the same setting as described above. For each agent, there was a small probability of (*p* = 0.02) per trial that the agent’s parameters will be re-sampled from the same uniform distribution. The overall number of trials in which the parameters were re-sampled was limited to four, and after each trial of parameters re-sampling the probability of another change was fixed to zero for 10 trials.

The combined loss function (Eq. 1) included the following *λ* hyperparameters values: 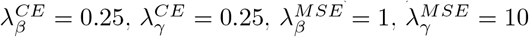, so that each loss term is approximately of equal magnitude.

#### d-RNN model

Similar to the one described above, with the two differences being are that the batch size was 200 and the LOOCV schema was performed over all 44 individuals.

#### Stationary hybrid exploration model

We adopted a similar model as described in [22], in which we estimated two stationary (i.e., fixed) parameters for each individual separately. Namely, a *β* random exploration parameter and a *γ* directed exploration parameter using maximum-likelihood estimation.

#### Bayesian particle filtering

For each individual, we set 1,000 particles to approximate the distribution of two hidden RL parameters {*β, γ*}. Initial distribution of the particles were set to be Gaussians, with means of {0, *−*2} and variances of {1, 1}, respectively. The hyper-prior for the dynamics of *β* was *σ*_*β*_ = 0.1, and for *γ* was *σ*_*γ*_ = 0.1. We clipped the *β* to range between (0,4) to match the estimation range of other models.

## Acknowledgments

We thank Yuval Ger for her help with the figure design. The study was funded by the Israel Ministry of Science and Technology, and the Israeli Science Foundation (Grant 2536/20 awarded to NS).

## Supporting Information

**S1 Table. Sensitivity of t-RNN to parameters quantization**. To ensure that quantization did not affect our performance, we ran two experiments where we quantized the RL parameters to as low as 3 bins (*α* and *β* parameters to 3 evenly spaced bins each) and up to 10 bins (*α* and *β* parameters to 10 evenly spaced bins each). We found similar performance across all three quantization settings.

**Table S1.** Action prediction (BCE) and parameters estimation (MSE) for different sizes of quantization bins of t-RNN. Averaged across *N* = 30 artificial test-agents. *↓* Lower is better. Mean *±* SD.

| Bins size | Action (BCE ↓) | $\alpha$ (MSE ↓) | $\beta$ (MSE ↓) |
| --- | --- | --- | --- |
| $3 \times 3$ | 0.321±0.12 | 0.024±0.01 | 0.020±0.01 |
| $5 \times 5$ (main text) | 0.321±0.12 | 0.024±0.01 | 0.019±0.01 |
| $10 \times 10$ | 0.322±0.13 | 0.025±0.01 | 0.019±0.01 |

**S2 Table. Action prediction and RL parameter recovery of simulated test agents**. Summary table of the raw results presented in the validation study of the main text (see Fig. 2A).

**Table S2.** Action prediction (BCE) and parameters estimation (MSE) of simulated data. Averaged across *N* = 30 artificial test-agents. *↓* Lower is better. Mean *±* SD.

| Model | Action (BCE ↓) | $\alpha$ (MSE ↓) | $\beta$ (MSE ↓) |
| --- | --- | --- | --- |
| Q-stationarity | 0.350 ± 0.15 | 0.053 ± 0.10 | 0.073 ± 0.10 |
| Bayesian (particle filter) | 0.327 ± 0.13 | 0.031 ± 0.03 | 0.038 ± 0.03 |
| t-RNN | 0.321 ± 0.12 | 0.024 ± 0.01 | 0.019 ± 0.01 |

**S3 Table. Volatile environment**. To assess the performance of t-RNN model in a volatile environment, we conducted an additional analysis similar to the one described in the main text (see Validating t-RNN using synthetic behavior) with agents simulated in a two-armed bandit task. However, here instead of a fixed reward probability for each arm, the reward expected value schedule was governed by a random walk with a drift rate of 0.025, and upper and lower bounds of 0.15 and 0.85, respectively. Our findings align with the conclusions mentioned in the main text, demonstrating that the t-RNN outperformed the alternatives in terms of both action prediction and parameter estimation. Therefore, we conclude that our conclusions regarding tRNN performance generalized to two-armed bandit tasks in a volatile environment.

**Table S3.**
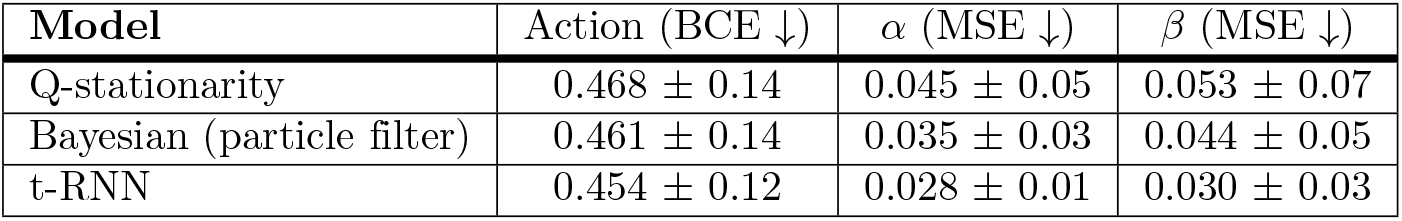
Action prediction (BCE) and parameters estimation (MSE) of simulated data in a volatile environment. Averaged across *N* = 30 artificial test agents. *↓* Lower is better. Mean *±* SD.

| Model | Action (BCE ↓) | $\alpha$ (MSE ↓) | $\beta$ (MSE ↓) |
| --- | --- | --- | --- |
| Q-stationarity | 0.468 ± 0.14 | 0.045 ± 0.05 | 0.053 ± 0.07 |
| Bayesian (particle filter) | 0.461 ± 0.14 | 0.035 ± 0.03 | 0.044 ± 0.05 |
| t-RNN | 0.454 ± 0.12 | 0.028 ± 0.01 | 0.030 ± 0.03 |

**S4. Sensitivity of t-RNN as a function of the number of observations and free parameters in the theoretical RL model**. Here, we examine the impact of the number of trials available and the number of free parameters for the theoretical RL parameters on the quality of t-RNN parameter recovery. Specifically, we compared the performance of the stationary maximum-likelihood (MLE; fitted with 5 different search locations) and the t-RNN models trained with one, two, or three theoretical free parameters. Using the Q-learning and preservation model, we systematically increased the number of free parameters (see Table Experiment specification.) and performed parameter recovery on 150 test agents (simulated with stable parameters). Additionally, we systematically increased the number of trials used to find optimal parameters in increments of 25. We computed the mean-square error (MSE) between the true parameters and the estimated parameters.

Results revealed that when the RL theoretical model included only one free parameter (i.e., learning rate) stationary MLE method showed better performance compared with t-RNN. However, when we included two or three free theoretical parameters, t-RNN did not fall behind the MLE approach. Importantly, we found preliminary results suggesting that for agents with more than one free parameter, t-RNN might outperform the MLE approach when the data has a very low number of observations per individual (*∼* 25 to 50). Furthermore, a close examination of the results shows that the MLE method for models with more than one parameter showed somewhat less consistent improvement in MSE as a function of trials compared with t-RNN (see Fig. S4). This is most likely the result of MLE being more sensitive to initial values (we used five starting points that were chosen randomly for each agent, and used for the results the starting values that gave the best result). Overall, t-RNN has the advantage of being able to predict dynamic parameters across trials, and the current analysis adds that when the thermotical parameters are stable, t-RNN does not fall behind conventional MLE approach and might even outperform MLE when the number of observations is limited.

**Table S4.** Experiment specification.

|  |  |  |  |
| --- | --- | --- | --- |
| Exp. 1 | $\alpha \sim U(0, 1)$ | $\beta = 3$ (fix) | $\kappa = 0$ (fix) |
| Exp. 2 | $\alpha \sim U(0, 1)$ | $\beta \sim U(0, 10)$ | $\kappa = 0$ (fix) |
| Exp. 3 | $\alpha \sim U(0, 1)$ | $\beta \sim U(0, 10)$ | $\kappa \sim U(-.5, .5)$ |

**Fig S4.**
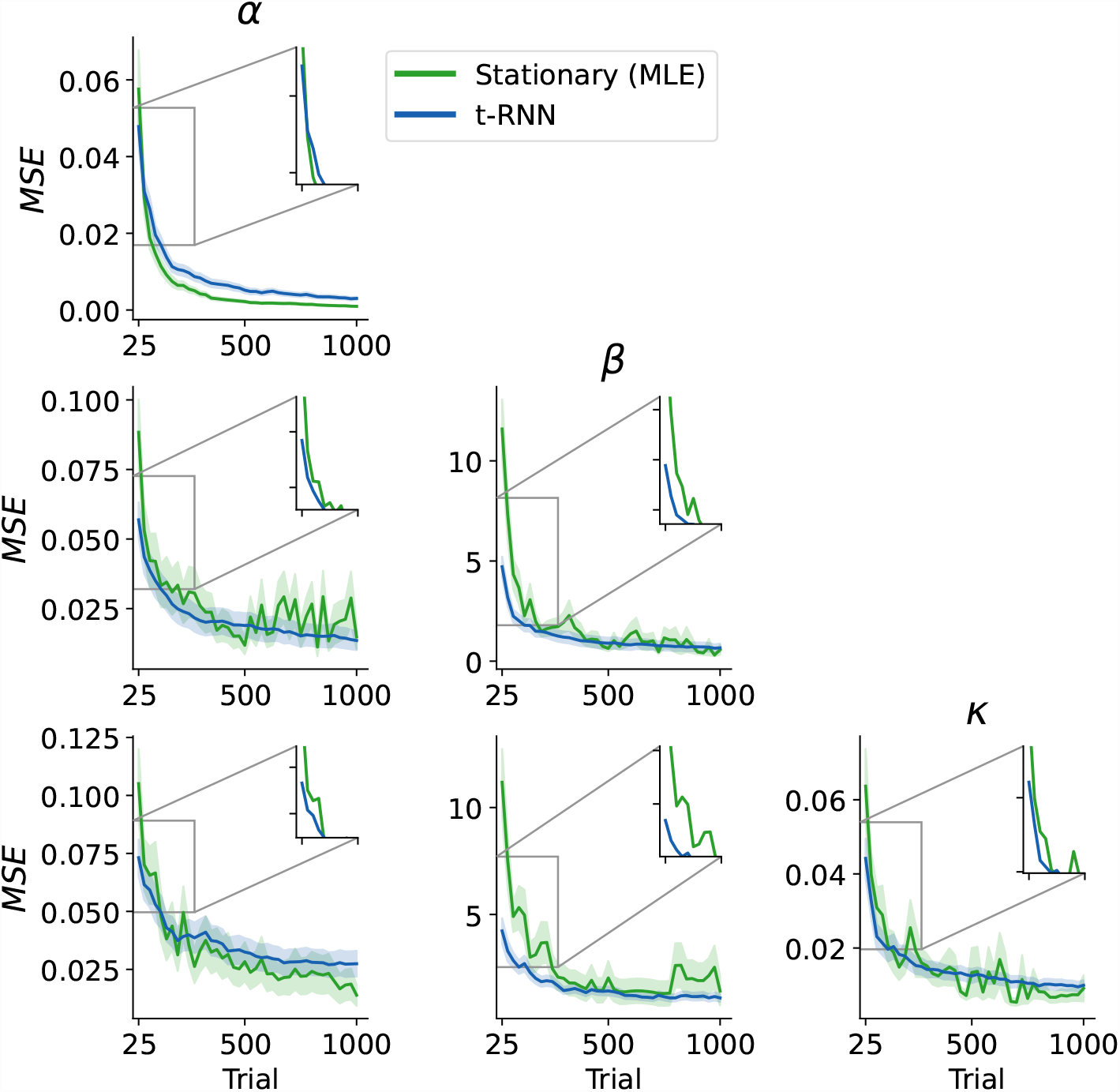
Sensitivity to the trial numbers and free parameters. Parameter recovery performance (measured in MSE and averaged over 150 test agents; y-axis) for the stationary (maximum-likelihood; green) and t-RNN (blue) as a function of available trials (x-axis). We also vary the number of free parameters, represented by: the top, middle, and bottom panels for Exp. 1, Exp. 2, and Exp. 3, respectively (shaded area signifies s.e.m; top right corner zoom in on first 100 trials).

**S5 Table. Action prediction behavioral dataset [9]**. Summary table of the raw results presented in the main text (see Fig. 3A).

**Table S5.**
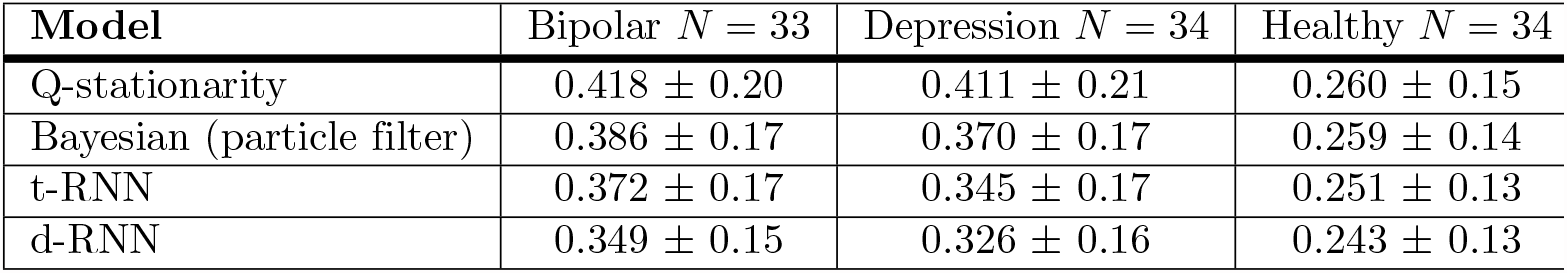
Action prediction for each model divided by diagnostic label. Behavioral dataset from [9]. Measured with binary cross-entropy (BCE; *↓* lower is better). Mean *±* SD.

**S6 Table. Summary statistics of RL parameters estimations by diagnostic group** Summary statistics of the trial-by-trial RL parameter estimation produced by t-RNN for each diagnostic group.

**Table S6.**
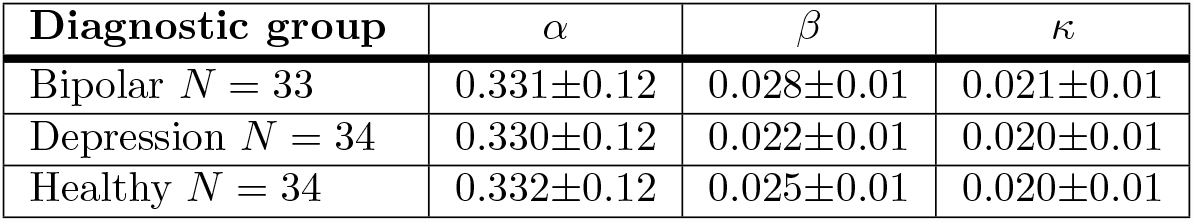
RL parameter estimation for each diagnostic group. Behavioral dataset from [9]. Mean *±* SD.

**S7. Fig. Relation of perseveration estimation with stay probability**. Further details examples of the analysis presented in the main text (see Fig 3C), in which we computed a Pearson correlation between the moving average stay probabilities and the t-RNN trial-by-trial *κ* perseveration estimation.

**Fig S7.**
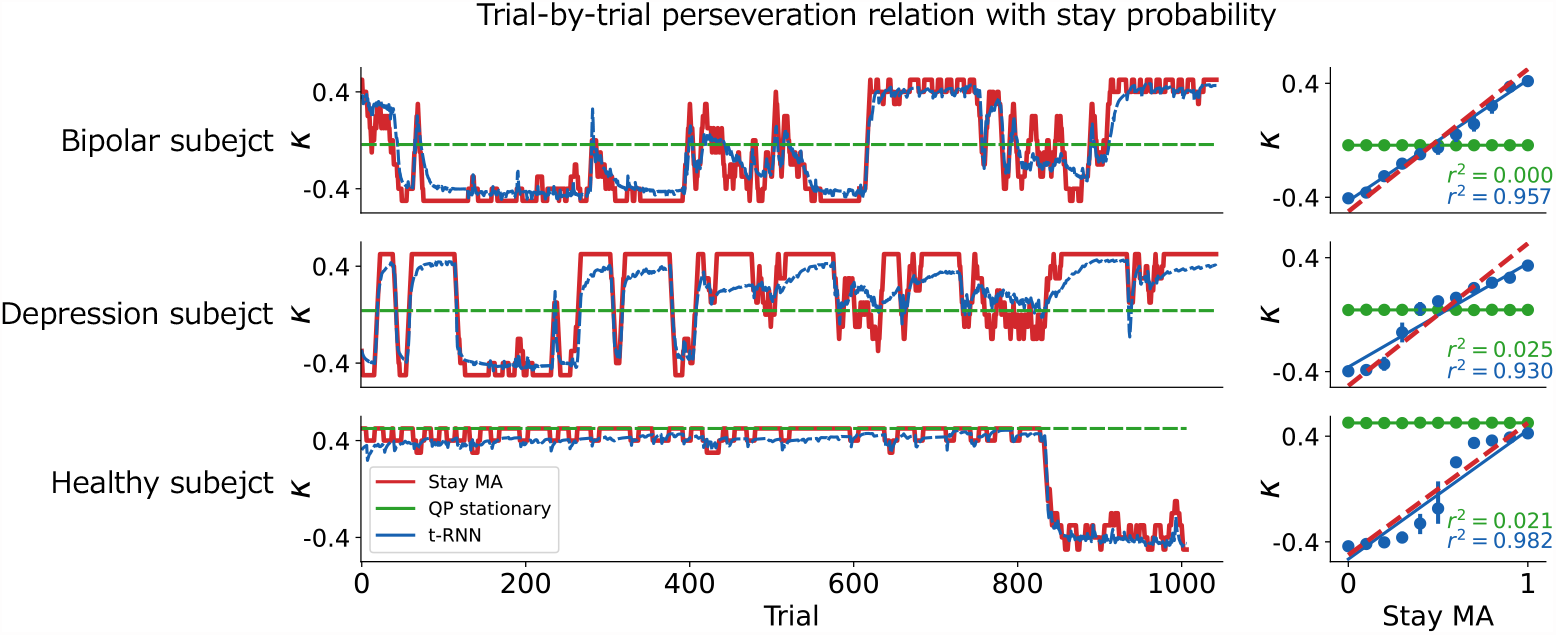
The left panels show the trial-by-trial RL *κ* perseveration parameter estimation using t-RNN (blue) and QP-stationarity (green) methods, along with the moving average calculation of the stay probabilities (red; window size of 10 trials) for three example subjects (one from each diagnostic group). The right panels show the corresponding Pearson correlation between the moving average stay probabilities and *κ* parameter estimation of t-RNN and QP-stationarity model (red dashed line denotes the identity). The results indicate a strong correlation between the stay probabilities and *κ* parameter estimation of t-RNN (*r*^2^ *>* 0.9), but not of the QP-stationarity (*r≈* 0), which fails to detect changes in subject behavior throughout the task.

**S8. Fig. Additional analysis inverse-temperature estimation**. In the main text, our analysis mainly focused on the dynamical *κ* perseveration parameter estimation produced by t-RNN. For completeness, we provide here a similar analysis for the *β* inverse-temperature which determines the randomness of action selection. To further support the interpretability of t-RNN parameters, in Fig S8C we used another easy-to-interpret model-agnostic measurement. We calculated a moving average of the distance between t-RNN action prediction probabilities and the probabilities of random choice policy (i.e., |*p*_*tRNN*_ (*a*_*t*_) *−* 0.5| ; window size of 10 trials). If t-RNN *β* estimates are related to the degree of randomness in action selection, we should expect a high correlation with the moving average model-agnostic estimate. We calculated a Pearson correlation between the moving average and t-RNN trial-by-trial *β* estimations for each individual. Overall, we found that the median Pearson correlation across individuals was very high (Median=0.74).

**Fig S8.**
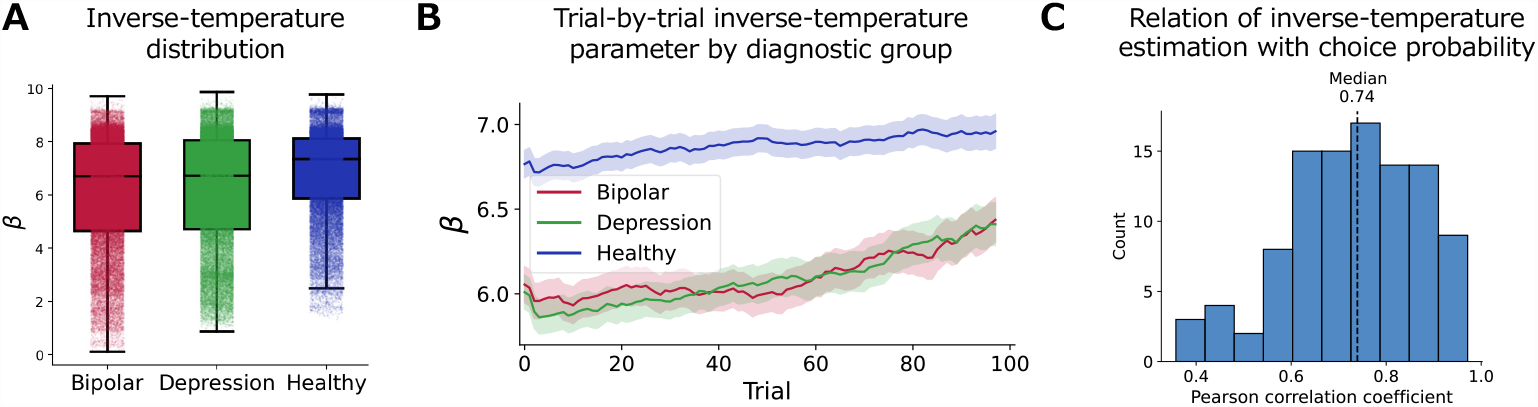
Analysis of the t-RNN trial-by-trial *β* inverse-temperature estimation. **(A)** Boxplot of the time-varying *β* estimates for each diagnostic group (middle black solid lines denote the median; light dots indicate a single trial estimate of the *κ* preservation parameter). Results indicate that all groups exhibit similar estimates, with a slight increase in the healthy group compared to the clinical groups. This suggests that the healthy group’s action selection was slightly less random than the clinical group’s. **(B)** Trial-by-trial *β* inverse-temperature parameter estimates by t-RNN averaged over the first 100 trials of each block and for each diagnostic group separately (shaded area signifies the s.e.m). In all three groups, subjects show an increase in their *β* estimates (behavior became less random as the block progressed), but the baseline differs, with clinical groups having a lower starting point. **(C)** Distribution of the Pearson correlation between the choice probabilities produced by t-RNN (calculated using a moving average of 10 trials) and the time-varying *β* inverse-temperature produced by t-RNN for each subject individually. The results indicate a strong relationship between t-RNN time-varying RL *β* parameter estimation and the moving average of the choice probabilities.

**S9. Fig. Relation of inverse-temperature estimation with random choice**. To illustrate our analysis approach, as shown in Fig S8C, we showcase the results of three example subjects, one from each diagnostic group.

**Fig S9.**
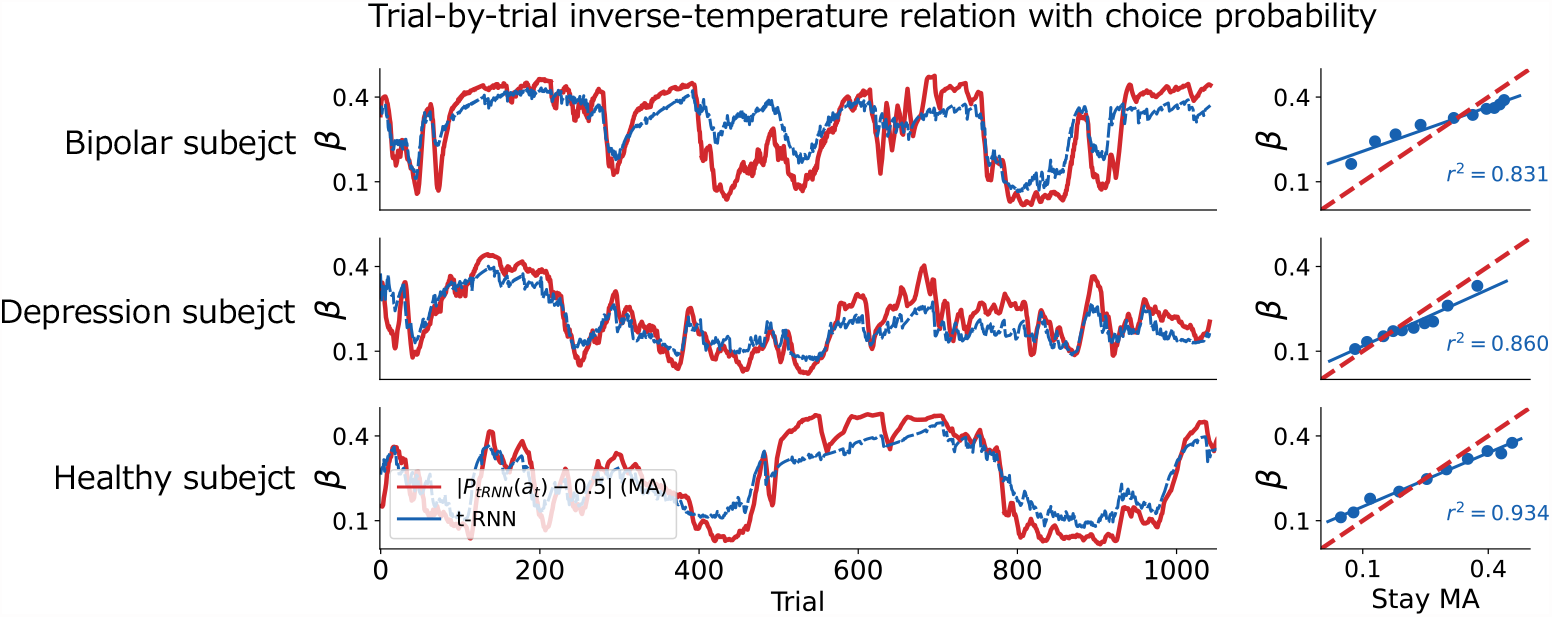
The left panels show the trial-by-trial RL *β* parameter estimation of t-RNN (blue; divided by 20 to fit the scale) and a moving average calculation of the absolute value difference between the choice probabilities produced by t-RNN and random choice probability (red; window size of 10 trials) for three example subjects (one from each diagnostic group). The right panels show the corresponding Pearson correlation between the moving average choice probabilities and *β* parameter estimation of t-RNN (red dashed line denotes the identity). The results indicate a strong correlation between the two estimations.

**S10. Fig. Supplement Fig. 3D**. We present here the full trial trajectories and model predictions of the two subjects presented in Fig. 3D.

**Fig S10.**
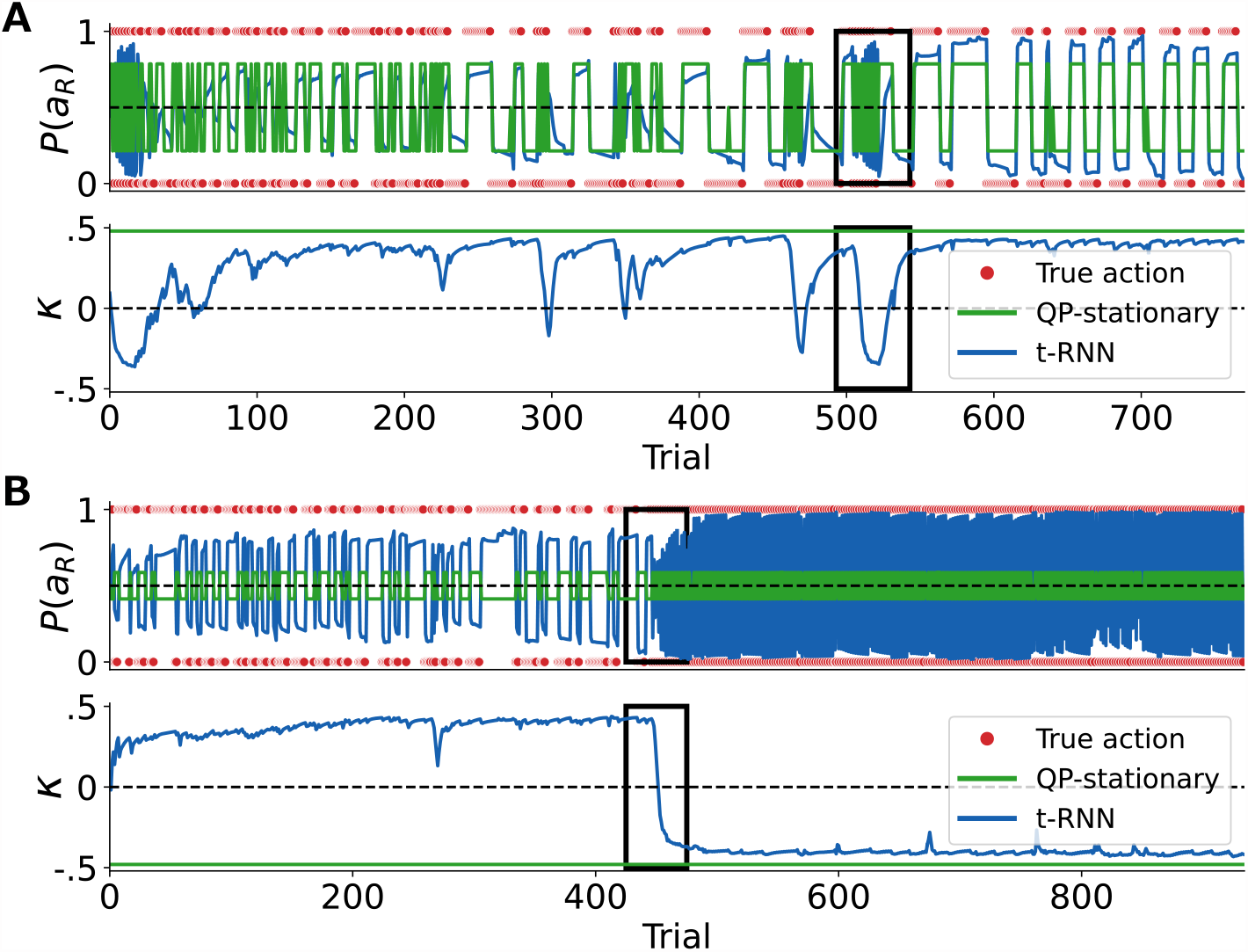
Supplement Fig. 3D. In Fig. 3D we depicted specific trials to illustrate the estimation of parameters across the two methods (QP-stationary, t-RNN). For completeness, we present the full range of trials for the same subjects (black boxes represent the trials that are depicted in Fig. 3D). **(A)** Example bipolar subject. **(B)** Example depression subject. Action prediction (top) and theoretical RL *κ* parameters estimation (bottom).

**S11 Table. Action prediction behavioral dataset [22]**. Summary table of the raw results presented in the main text (see Fig. 4A).

**Table S11.** Action prediction for each model. Behavioral dataset from [22]. Measured with binary cross-entropy. *↓* lower is better. *N* = 44. Mean *±* SD.

| Model | BCE ↓ |
| --- | --- |
| Stationary hybrid explore. | $0.331 \pm 0.07$ |
| Bayesian (particle filter) | $0.332 \pm 0.07$ |
| t-RNN | $0.320 \pm 0.07$ |
| d-RNN | $0.310 \pm 0.08$ |

**S12. Fig. Directed exploration dynamics of repeated action blocks**. We conducted an additional investigation into the trial-by-trial dynamics of the t-RNN theoretical parameter estimation of the behavioral dataset [22]. Specifically, we identified a block of 10 consecutive trials where an individual persisted in choosing the same option throughout. Consistent with our expectations, we observed a significant reduction in the t-RNN’s *γ* parameter estimation of directed exploration, indicating a lack of interest in exploring the less familiar option (see Fig. S12A). To ascertain whether this pattern holds true across individuals and trials, we aggregated the t-RNN’s *γ* parameter estimates over all blocks (a total of 70 blocks) where individuals chose the same option. Our findings revealed a similar trend of a sharp decrease in the *γ* directed exploration parameter estimation (mean*±*s.e.m. before: 0.52*±*.02, after: 0.18*±*0.01).

**Fig S12.**
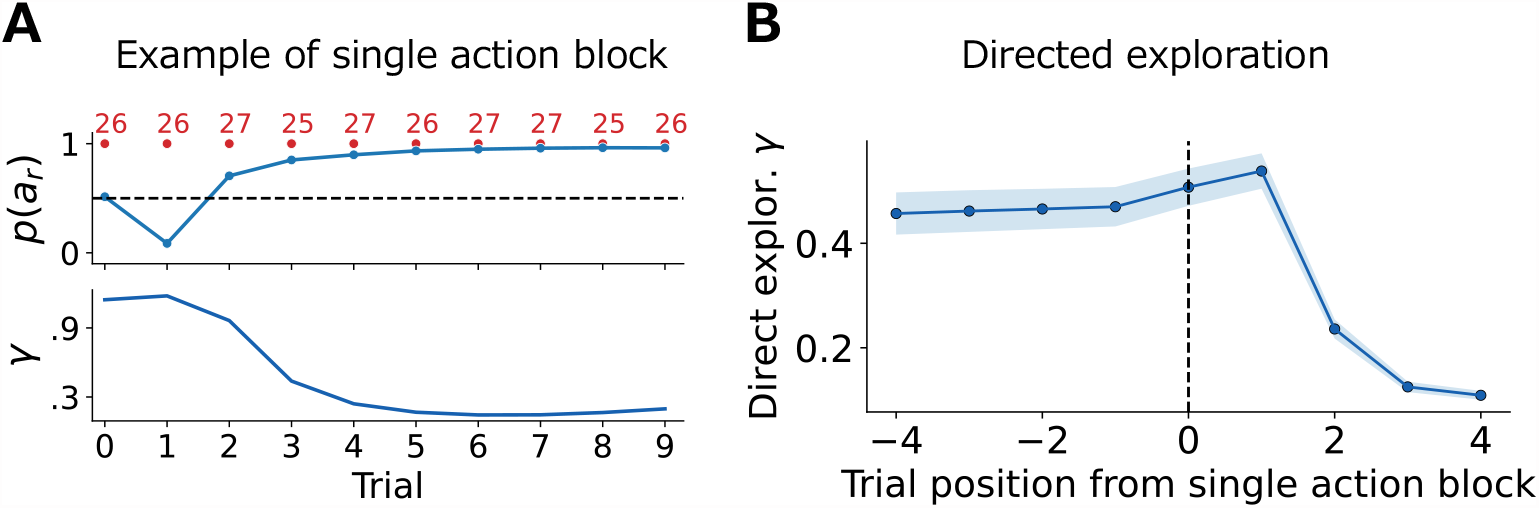
**(A)** Example of a single instantiation of a single action block. The top panel depicts the sequence of chosen actions, obtain rewards, and t-RNN action predictions as red dots, red numbers, and blue solid lines, respectively. The subject consecutively chooses the same action. The bottom panel depicts t-RNN *γ* directed exploration parameter estimation, where it estimates a step decrease in the parameter throughout the block. **(B)** Dynamics of the directed exploration parameter averaged across all instantiations of a single action block. t-RNN estimates a sharp decrease in the *γ* directed exploration parameter in these blocks.

**S13. Fig. t-RNN concatenation ablation**. We conducted an ablation analysis in which we trained a t-RNN model without concatenation of the categorical class predictions of the RL parameters into the action prediction head, and compared it to the original method that was trained with concatenation. Results revealed that training t-RNN without concatenation led to a degradation in the performance in the action prediction for both empirical datasets tested (Wilcoxon signed rank test; *** *p <* .001 for Dezfouli et al. dataset [9]; ** *p <* .01 for Gershman dataset [22]).

**Fig S13.**
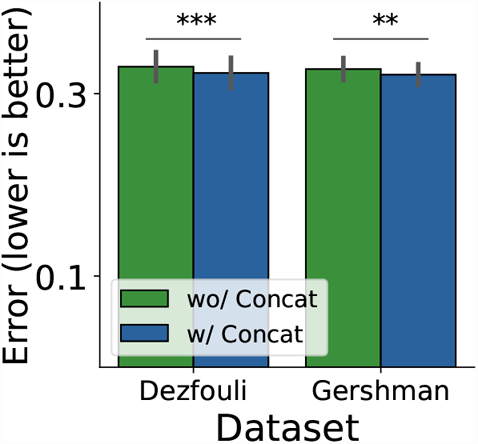
t-RNN concatenation ablation. Training t-RNN without concatenation (green bar) led to a degradation in the performance in the action prediction compared to t-RNN trained with concatenation (blue bar) for both empirical datasets (error measured in BCE; black lines indicate s.e.m; *** *p <* .001; ** *p <* .01)

